# Altered beta-band oscillations and connectivity underlie detail-oriented visual processing in autism

**DOI:** 10.1101/2020.01.30.926261

**Authors:** Luca Ronconi, Andrea Vitale, Alessandra Federici, Elisa Pini, Massimo Molteni, Luca Casartelli

## Abstract

Sensory and perceptual anomalies have been increasingly recognized as core phenotypic markers for autism spectrum disorders (ASD). A neurophysiological characterization of these anomalies is of utmost importance to understand more complex behavioural manifestations within the spectrum. The present study employed electroencephalography (EEG) to test whether detail-oriented visual perception, a recognized hallmark of ASD, is associated with altered neural oscillations and functional connectivity in beta (and alpha) frequency bands, considering their role in feedback and top-down reentrant signalling in the typical population. A sample of children with diagnosis of ASD (N=18) and typically developing peers (TD; N=20) performed a visual crowding task, where they had to discriminate a peripheral target letter surrounded by flankers at different distances, together with a control condition with no flankers. In TD participants the amplitude of the target-locked N1 component and its cortical sources was significantly modulated as a function of visual crowding, whereas such modulation was absent in the ASD group, suggesting that their visual scene analysis takes place without a flexible neural computation. The comparison between groups showed a decreased activity in the ASD group in occipital, infero-temporal and inferior/middle frontal regions in conditions requiring detail-oriented perception as opposed to conditions requiring the discrimination of target in isolation. Moreover, in TD participants detail-oriented perception was associated with an event-related beta power reduction (15-30 Hz), which was not evident in the ASD group. A data-driven functional connectivity analysis highlighted in the ASD sample an increased connectivity in the beta frequency range between occipital and infero-temporal regions. Notably, individual hyperconnectivity indexes correlated to less severe ASD symptomatology and to a diminished detail-oriented perception, suggesting a potential compensatory mechanism. Overall, these results show that altered communication in the beta frequency band may explain atypical perception in ASD, reflecting aberrant feedback connectivity within the visual system with potential cascade effects in visual scene parsing and higher-order functions.

## Introduction

Autism spectrum disorder (ASD) is a severe neurodevelopmental condition affecting ∼1% of the population and causing impairments in social communication and interaction, as well as restricted interests and activities (Association, 2013). A consistent body of evidence in the last two decades has associated ASD with anomalies in perception and sensory processing (for reviews see: Dakin and Frith, 2005; Simmons *et al*., 2009; Pellicano and Burr, 2012; Ronconi *et al*., 2016; Robertson and Baron-Cohen, 2017). This has gradually determined a partial change of perspective in the field of autism research. While recognizing the importance of the core social, communicative and cognitive difficulties associated with the condition, the need of a better understanding of low-level sensory and perceptual anomalies has acquired increasing importance. Accordingly, a recent revision of the diagnostic criteria for ASD has brought sensory processing as a key domain of the disorder (Association, 2013). A paradigm shift that focuses more on basic sensory processing and its association with dysregulated neural activity could benefit ASD research for different reasons (Vissers *et al*., 2012). First, to improve early detection, considering that impairments or anomalies in basic sensory functions and their neural underpinnings can be used as markers of ASD. Furthermore, (reh)abilitation protocols could benefit from a better awareness of the different sensory profiles. Second, to support translational research, given that the identification of core sensory and oscillatory anomalies is easier to evaluate in animal models and easier to map onto specific genetic/epigenetic factors as compared to more complex constructs related to social cognition. Finally, basic sensory and cognitive anomalies may help to characterize the ontogeny of social cognition, deconstructing them in more elementary components.

Recently, there has been an increasing awareness, not only in the context of ASD but also for other neuropsychiatric and neurodevelopmental disorders, about the relevance of describing the precise neural temporal dynamics associated to specific behavioural and cognitive phenotypes (Buzsaki and Watson, 2012; Uhlhaas and Singer, 2012). In particular, neural oscillations constitutes a fundamental mechanism for neural communication between different brain regions (Fries, 2005, 2015). A mechanistic approach that evaluates anomalies in oscillatory synchronization across both short- and large-scale cortical networks may be extremely useful in understanding some core manifestations of ASD (Kessler *et al*., 2016; Seymour *et al*., 2019), in line with several accounts claiming that ASD is primarily a disorder of brain connectivity (Belmonte *et al*., 2004; Wass, 2011; Vissers *et al*., 2012).

In the domain of vision, information processing is considered markedly atypical in ASD. Indeed, individuals with ASD show higher performance in detail-oriented tasks (Dakin and Frith, 2005; Simmons *et al*., 2009; Robertson and Baron-Cohen, 2017), such as for example better visual search performance (O’Riordan *et al*., 2001; Manjaly *et al*., 2007; Joseph *et al*., 2009; Gliga *et al*., 2015), and better resilience to visual illusions (for reviews see Happé and Frith, 2006; Gori *et al*., 2016). However, individuals with ASD exhibit also an increased interference from irrelevant stimuli (Burack, 1994; Adams and Jarrold, 2012; Ronconi *et al*., 2013, 2018) and visual sensory overload has been well documented both at the behavioral and neurophysiological level (Pritchard *et al*., 1987; Belmonte, 2000; Kern *et al*., 2006; Leekam *et al*., 2007).

An effective probe to test the putative superiority in local visual information processing and distractor suppression is visual crowding, a perceptual phenomenon that is typically observed in peripheral vision consisting in a decreased ability to discriminate objects when presented with nearby flankers (Bouma, 1970). Crowding limits the recognition, not the detection, of visual stimuli of different complexity, ranging from simple objects such as oriented gratings, to more complex objects such as letters and faces (Levi, 2008; Pelli, 2008; Whitney and Levi, 2011). In the context of ASD, there have been behavioural reports of superior performance during visual crowding tasks, i.e. less interference from flankers (Baldassi *et al*., 2009; Keita *et al*., 2010), but also other studies showing a similar magnitude and spatial extent of crowding in individuals with ASD as compared to controls (Constable, 2010; Freyberg *et al*., 2016).

The neural computations involved in visual crowding seems to be dependent from the complexity of objects involved; i.e. the more complex the object, the higher the visual area involved in crowding. This might be partially responsible also for the mixed behavioural findings in ASD. For example, for simple stimuli such as oriented grating an excessive integration may occur predominantly at early stages of visual processing, like V1 and V2, where binding of elementary features occurs (Millin *et al*., 2013; Chen *et al*., 2014). For more complex objects, like letters or faces, crowding may predominantly occur at a higher level along the visual hierarchy, for example in the ventral stream areas like V4 and infero-temporal areas that mediates integration of object contours (Motter, 2006; Liu *et al*., 2009; Anderson *et al*., 2012). Moreover, resolving crowding for complex visual objects involves, like any other computation on complex visual inputs, top-down reentrant feedback loops from higher-order to lower-order visual areas, leading to the emergence of significant variations in the rhythmic oscillatory activity (Donner and Siegel, 2011; Siegel *et al*., 2012; Bastos *et al*., 2015; Jensen *et al*., 2015). Finally, independently from the complexity of objects, it has been shown that also dorsal-to-ventral feedback mediates segmentation of the target from the flankers through the activation of receptive fields of appropriate size (Lee *et al*., 1998; Vidyasagar, 1999, 2004; Lamme and Roelfsema, 2000), which also leads to variations in the rhythmic oscillatory activity (Vidyasagar, 2019).

Several electroencephalography (EEG) studies on visual crowding have been recently performed in the typical population using complex stimuli configurations (e.g. letters, Vernier stimuli). These studies provide a consistent picture of the neurophysiological correlates of crowding for complex objects, constituting an important starting point for better understanding enhanced local information processing in individuals with ASD. Indeed, these studies consistently showed that crowding induced a suppression in amplitude of the N1 event-related potential (ERP), peaking around 200/250 ms post-stimulus (Chicherov *et al*., 2014, Ronconi *et al*., 2016*a*). In terms of oscillatory correlates, crowding have been associated with neural oscillations in the beta band (15-30 Hz) (Ronconi *et al*., 2016*a*; Ronconi and Marotti, 2017; Battaglini *et al*., 2019). Particularly, Ronconi et al. (2016) measured crowding with different target-to-flankers spacing and found an event-related power reduction in the beta band that was enhanced in a strong crowding regime (i.e. smaller target-to-flankers distance) relative to a weak crowding regime (i.e. larger target-to-flankers distance). This beta power reduction also correlated with task performance at an individual level. Ronconi and Bellacosa Marotti (2017) further confirmed the selective relationship between beta band oscillations and visual crowding, by showing that beta power in the pre-stimulus time window was increased in frontal and parieto-occipital sensors for trials where participants correctly discriminated the target letter among flankers.

Neurophysiological studies in macaques and eletro-/magneto-encephalography (M/EEG) evidence in humans strongly suggest that oscillations in the alpha and beta-band mediate top-down feedback connectivity between the distant brain regions, e.g. between higher- and lower-order visual areas or between fronto-parietal and visual areas, constituting a fundamental mechanism for top-down processing in the visual domain (Donner and Siegel, 2011; Bastos *et al*., 2015; Jensen *et al*., 2015; Michalareas *et al*., 2016). This idea is in line with studies showing a link between beta oscillations and visual crowding reviewed above and, more generally, is in line with studies linking beta oscillations to other functional aspects involved in human vision like, such as spatial orienting of attention (Siegel *et al*., 2008; Buschman and Miller, 2009; Fiebelkorn *et al*., 2018), coherent motion discrimination (Aissani *et al*., 2014) and Gestalt perception (Zaretskaya and Bartels, 2015). Interestingly, these visual and attentional mechanisms have all been reported to be anomalous in ASD in previous behavioral studies (for reviews see Dakin and Frith, 2005; Simmons *et al*., 2009; Keehn *et al*., 2013, Ronconi *et al*., 2016; Robertson and Baron-Cohen, 2017). Overall, these results suggest that beta oscillations might be a relevant target for neurophysiological studies testing altered neural oscillations during visual processing in ASD, similarly to what have been recently shown for gamma band oscillations (Sun *et al*., 2012; Peiker *et al*., 2015) and alpha-gamma coupling (Seymour *et al*., 2019).

Taking advantage of the evidence accumulated so far on the role of beta band oscillations in visual and attentional tasks, including crowding, in the present study we employed high-density electroencephalography (EEG) in participants with ASD to investigate the neurophysiological (ERPs) and oscillatory correlates of local visual processing within a crowding regime, and to study the underlying network activity and connectivity pattern at the level of cortical sources. In particular, testing alpha/beta oscillations during local visual processing in ASD could inform about possible alterations in neural networks promoting top-down reentrant feedback connectivity coming from higher-order visual areas and from fronto-parietal attentional areas.

## METHODS

### Participants

Twenty-two children with ASD were recruited from the Scientific Institute IRCCS “E. Medea” (Bosisio Parini, Italy), and twenty-two typically developing (TD) children were recruited as control group from local schools in the same geographic area.

Participants with ASD were selected according to the following criteria: (i) full-scale IQ>70 as measured by the WISC-III or IV (Wechsler, 1993); (ii) normal/corrected-to-normal vision and normal hearing; (iii) absence of epilepsy and gross behavioural problems; (iv) absence of drug therapy; and (v) absence of developmental dyslexia or attention deficit hyperactivity disorder (Association, 2013).

All children with ASD were diagnosed by licensed clinicians in according to the DSM-IV criteria and to the Autism Diagnostic Observation Scale (Lord *et al*., 2002). Children of the TD group did not have a prior history of neurological and/or psychiatric disorders, and their cognitive level were assessed using two subtests of the WISC: Block design for performance and Vocabulary to test verbal abilities (see Table 1). The two groups were matched for chronological age (t_(36)_= −1.957, *p*= 0.058) and cognitive level (performance subtest: t_(36)_= 0.188, *p*= 0.852; see Table 1).

**Table 1.**
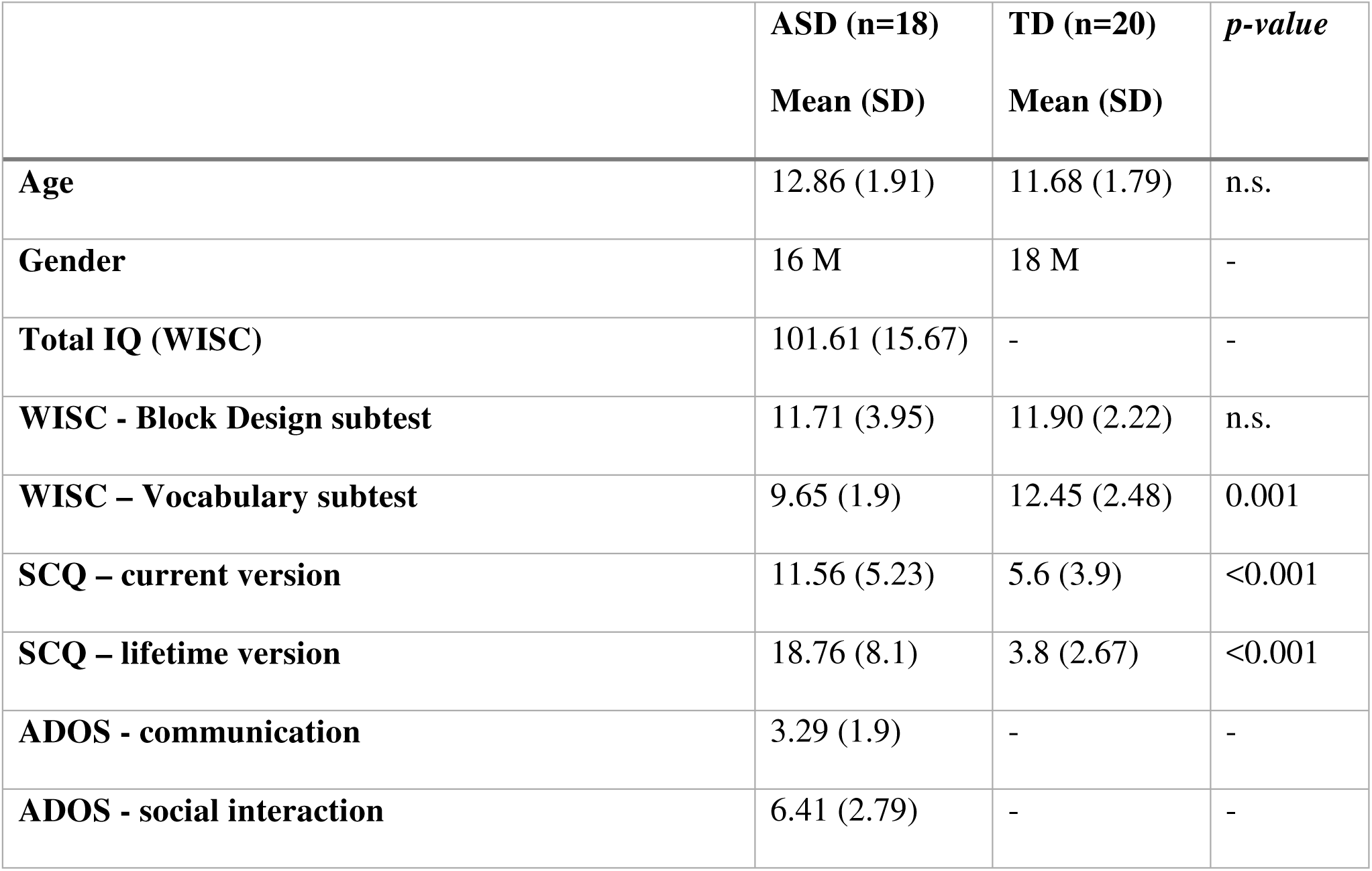
Descriptive statistics of participants (ASD=autism spectrum disorder; TD=typically developing; WISC=Wechsler Intelligence Scale for Children; SCQ= Social Communication Questionnaire; ADOS=Autism Diagnostic Observation Scale).

|  | **ASD (n=18)**<br>**Mean (SD)** | **TD (n=20)**<br>**Mean (SD)** | ***p-value*** |
| --- | --- | --- | --- |
| **Age** | 12.86 (1.91) | 11.68 (1.79) | n.s. |
| **Gender** | 16 M | 18 M | - |
| **Total IQ (WISC)** | 101.61 (15.67) | - | - |
| **WISC - Block Design subtest** | 11.71 (3.95) | 11.90 (2.22) | n.s. |
| **WISC – Vocabulary subtest** | 9.65 (1.9) | 12.45 (2.48) | 0.001 |
| **SCQ – current version** | 11.56 (5.23) | 5.6 (3.9) | <0.001 |
| **SCQ – lifetime version** | 18.76 (8.1) | 3.8 (2.67) | <0.001 |
| **ADOS - communication** | 3.29 (1.9) | - | - |
| **ADOS - social interaction** | 6.41 (2.79) | - | - |

One child belonging to the ASD group and one to the TD group were not able to complete the task behaviourally and therefore were excluded. Moreover, three children of the ASD group and one child of the TD group were excluded from the analysis because their EEG data were excessively contaminated by artefacts (e.g. too many eye movements during the stimulus presentation). Thus, the final sample comprised 18 children for the ASD group and 20 children for the TD group.

Informed consent was obtained from each child and their parents or legal tutor and the entire research protocol was conducted in accordance to the principles elucidated in the declaration of Helsinki of 2013. The ethical committee of Scientific Institute “E. Medea” approved the present study protocol.

### Apparatus and stimuli

Participants were seated in a dimly and quite room in front of a Philips 19S L LCD screen (19in., 75Hz) at 50cm. Stimulus presentation and data acquisition were performed with E-Prime2 (Psychology Software Tools, Inc. www.pstnet.com).

The stimuli consisted in 1.5 × 1.5 deg gaborized T-like, H-like or random configurations presented at full contrast (Michelson) on a mid-level grey background. Stimuli were created with a matrix of the size of the stimulus divided into a 5 × 5 grid of equally spaced x, y locations, using Psychtoolbox for Matlab (Brainard, 1997) (for details see Ronconi *et al*., 2016*a*).

The two letter stimuli (T and H) were created with the Gabors placed according to the letter configuration. Ts and Hs were composed, respectively, by 9 and 13 patches that could be both horizontal and vertical, and a centre-to-centre distance between adjacent patches was kept constant at 0.3 deg.

For the random configurations (i.e., fillers), the Gabors were filled in thirteen random location within the grid. Six random configurations were selected, and they were then rotated on plane of 90, 180 and 270 deg. Thus, we ended up with 24 different fillers to use in our final stimuli. Indeed, the final displayed stimuli were composed by seven configurations containing letters or fillers, vertically arranged at 1.9 deg centre-to-centre distance (Fig. 1). Overall the final stimulus covered an area of 1.5 × 13.3 deg on the display.

**Figure 1.**
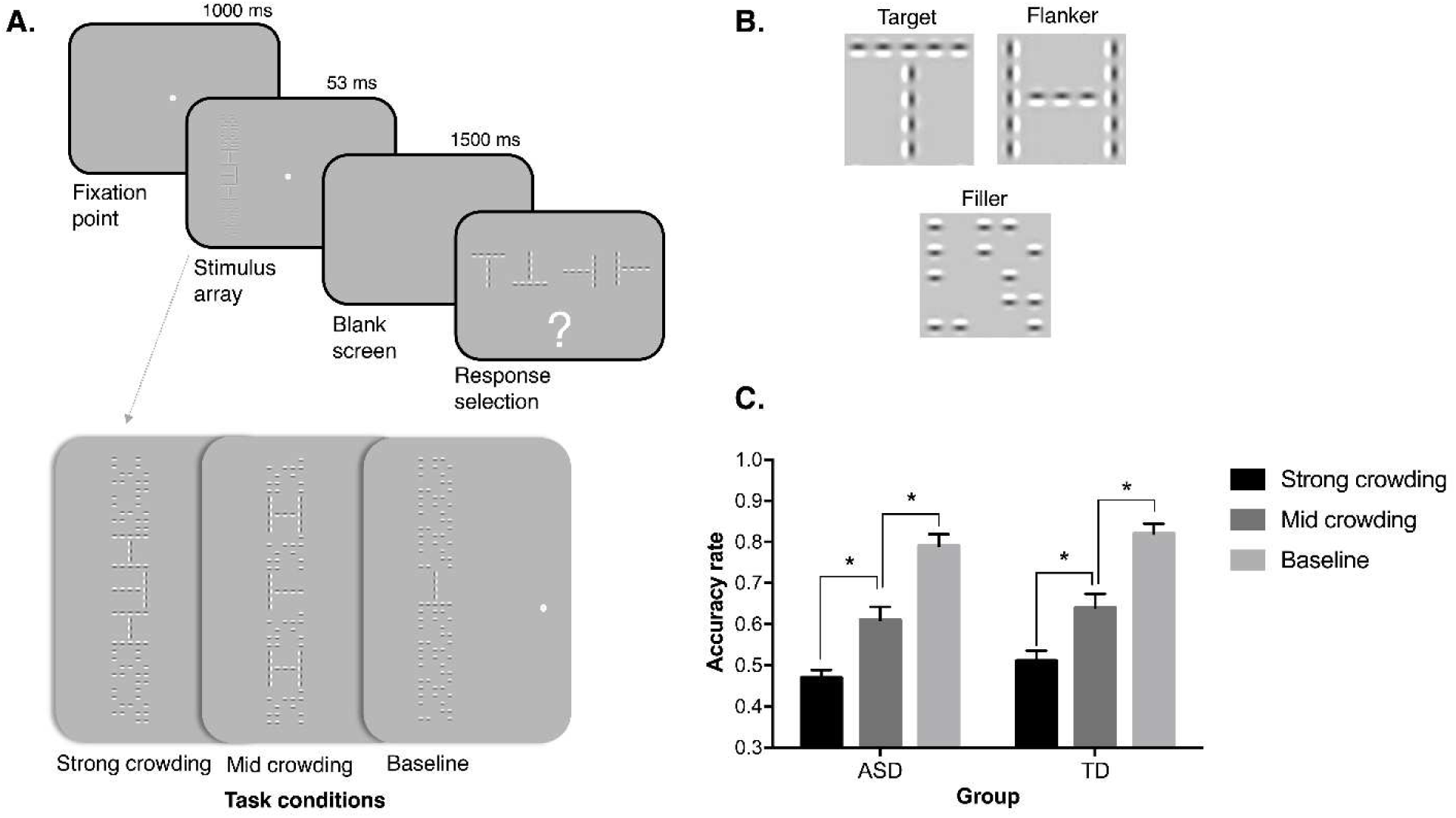
Schematic representation of the task procedure. (A) Example of a trial of the visual crowding task employed in the present study, where participants with ASD and TD peers were asked to discriminate the orientation of the peripheral target letter among the four possible alternatives. There were three task conditions, labelled as *strong* and *mid* crowding when flankers appear nearby the target at a closer or farther distance, respectively, and a *baseline* condition where the target was displayed in isolation. (B) Three examples of stimuli employed in each trial; fillers were used to ensure a constant visual stimulation across the different task conditions. (D) Behavioural results (accuracy rates) as a function of the task condition (error bars represent SEM; * = p<.05).

The final stimulus has the following arrangement: a target (T) in the centre position vertically flanked by six irrelevant configurations (i.e., flankers and/or fillers), three above and three below. The flankers (Hs) could be displayed either near the target T or in the intermediate position, and all the fillers occupied the locations left over. Both T and H could have one of four possible rotations (0– 270 deg in step of 90 deg) selected independently among them and randomly at each trial.

The arrangement of the Hs was selected in order to create the three experimental conditions (Figure 1). In the *strong* crowding condition, the Hs were placed nearby above and below the target and the remaining four positions were occupied by fillers. In the *mid* crowding condition, the Hs were placed in the intermediate positions, whereas fillers were displayed nearby and far from the T, both above and below. In the *baseline* conditions, fillers occupied all the locations and only the target letter T was displayed. These random configuration (i.e., fillers) were created in order to maintain a constant visual stimulation across the different crowding conditions and to ensure the same external masking for flanker letters (Hs) in both the *strong* and the *mid* crowding condition (for details see Ronconi *et al*., 2016*a*).

### Procedure

Participants performed an orientation discrimination task, where they were asked to identify which was the orientation of the central target letter (T). On each trial, a black fixation cross was centrally displayed for 1 s. The stimulus configuration was then briefly flashed (53 ms) at 11 deg eccentricity, half of the time to the left and half to the right of the fixation cross. Note that the chosen duration of stimuli presentation was below the time needed to execute a saccade towards the stimulus. A blank screen was then shown for 1.5 s. Finally, we presented a response display showing the four possible T rotations (figure 2). An inter trial interval of 500 ms containing a blank screen was used to avoid interference of motor artefact within the pre-stimulus temporal window. Participants were asked to keep their gaze fixed on the cross for the entire trial duration and then to communicate the orientation of the target to the experimenter who entered the response on the keyboard. We presented 528 trials in total, 176 trials for each of the three crowding conditions, randomly and equally distributed between left and right hemifields and among the four different orientations of the T. The entire session was divided in eight blocks in order to prevent fatigue. Before starting the acquisition, participants were presented with few practice trials (n=14) in order to become familiar with the task.

**Figure 2.**
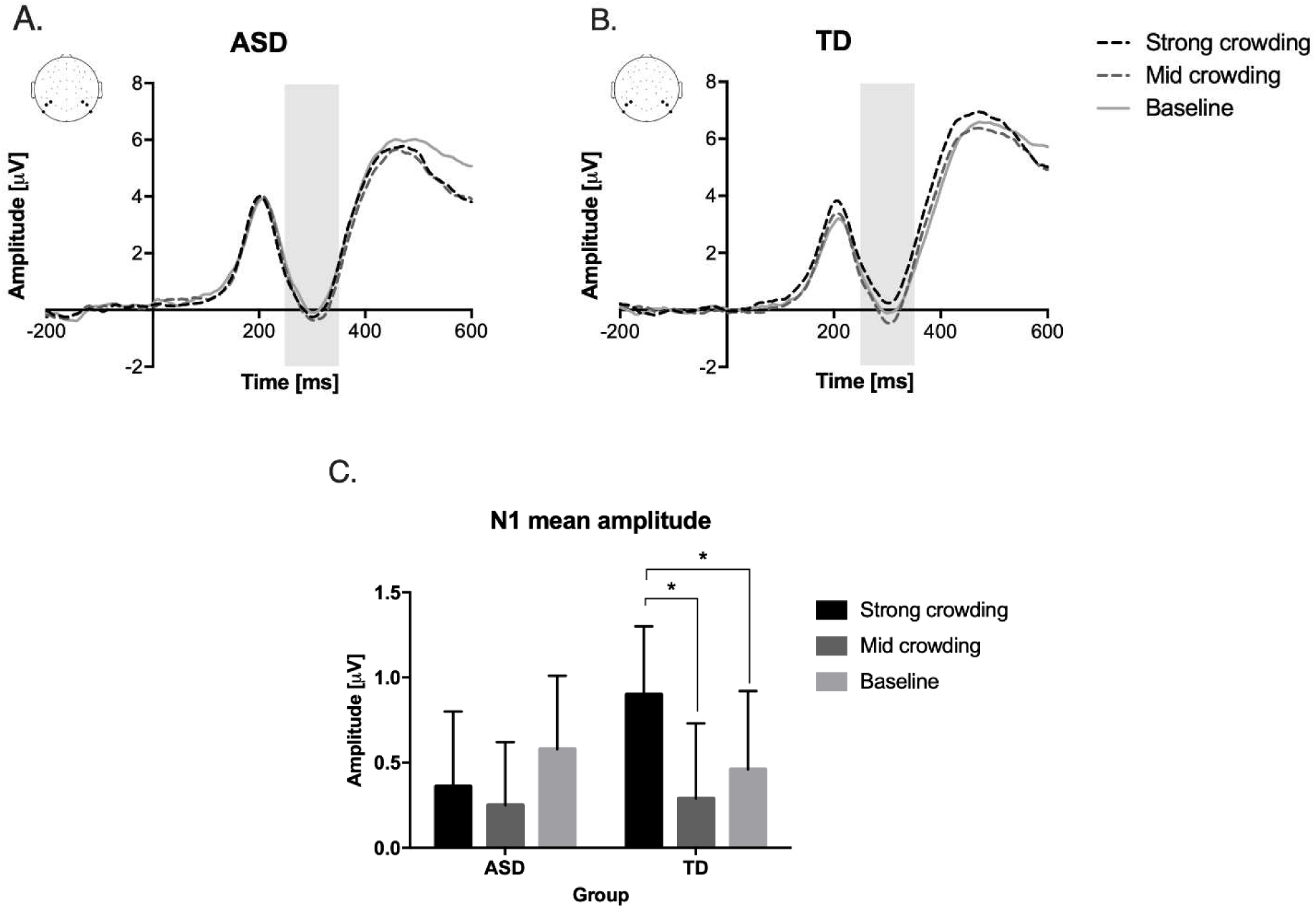
Event-related potential (ERPs) results. (A, B) Target evoked ERPs as a function of task conditions in the ASD and TD groups; data were obtained by averaging the activity in the posterior electrodes of interest (see upper-left insets). Activity in the grey shaded window in the time period 150-250 ms (N1) relative to the target onset were found to be differently modulated in the two groups as a function of task conditions. (C) Mean amplitude of this N1 ERP component as a function of group and task condition, revealing that while visual crowding modulated the amplitude of this component in the TD group, this modulation was absent in the ASD group (error bars represent SEM; * = p<.05).

### EEG recording and pre-processing

While children performed the task, their neural activity was recorded by using an Electrical Geodesics EEG system with 64-channel (Hydrocel Geodesic Sensor Nets, Electrical Geodesics, Inc.).

Input data were analog-filtered between 0.01 and 100 Hz and the sampling rate was 500 Hz. Offline, data were downsampled at 250 Hz and the reference were recomputed to the average. Then, data were filtered with a notch-filter at 50 Hz (non-causal Parks-McClellan Notch, order=180) and band-pass filtered between 0.1 and 80 Hz (non-causal IIR, order=2). The pre-processing steps, including filtering, artifact correction/removal and epoching, were performed in MATLAB using EEGLAB functions (Delorme and Makeig, 2004). The cleaned data segments were imported in Brainstorm (Tadel *et al*., 2011) to further estimate the location of active neural sources and functional connectivity, in addition to custom scripts.

### Data analysis - Behavioural data

For each participant we extracted the accuracy rate in each of the three crowding conditions. On this measure, we performed a repeated measure ANOVA with level of crowding (*strong*, *mid* and *baseline*) as within-subjects factor and group (ASD vs. TD) as between-subjects factor.

### Data analysis - Event-related potential (ERPs)

EEG epochs initially extracted for the analysis ranged between –1500 and 1500 ms relative to the target onset. Bad channels were interpolated when required (mean percentage ± SD: 4.9 ± 1.2 for the TD children and 3.8 ± 1.6 for the ASD group).

Epochs containing voltage deviation exceeding ±150 µV between –700 and 700 ms were removed. Also, epochs containing massive muscular artefacts (i.e. high-frequency activity affecting the majority of channels and time points) as well as blinks and eye movements occurring right before the target onset and during the target presentation were removed. The remaining blinks and eye movements were corrected through an independent component analysis decomposition (ICA). After the entire process of artefact rejection, the 80% of trials for TD and 78% for ASD group were retained and further analysed.

ERP components of interest were identified based on the previous literature and their latencies were chosen through visual inspection of the waveforms. In particular, we analysed the P1 component, with a positive peak around 200 ms relative to the target onset, and the N1 component, with a negative peak around 300 ms relative to the target onset (Figure 2A, 2B). The N1 has been previously associated with visual crowding (Chicherov *et al*., 2014, Ronconi *et al*., 2016*a*). As shown by previous studies (Di Russo *et al*., 2002; Chicherov *et al*., 2014), the N1 originates mainly from cortical sources in the parietal cortex. Thus, according to this previous literature and to the visual inspection of the data, we selected for the ERP analysis two clusters of parietal channels both on the left (channels 27, 28, 30) and on the right hemisphere (channels 42, 44, 45).

In order to explore the effect of crowding in the two groups we performed an ANOVA both on the P1 mean amplitude (150 and 250 ms) and on the N1 mean amplitude (250 and 350 ms) with level of crowding (*strong*, *mid* and *baseline*) as a within-subjects factor and group (*ASD* vs. *TD*) as a between-subjects factor. We also tested planned comparisons in case of a significant interaction.

As an additional analysis, we decided to subtract from the ERPs mean amplitude of the *strong* and *mid* crowding conditions the mean amplitude of the *baseline* condition. This was done in order to investigate if there was a difference between groups specifically due to the effect of crowding, ruling out the impact of the activity evoked purely by the processing of an isolated stimulus (i.e. *baseline* condition). On these measures we performed a repeated measure ANOVA with two levels of crowding (*strong-baseline* vs. *mid-baseline*) as within-subjects factor and group as between-subjects factor.

### Data analysis - Time-frequency decomposition

For the analysis of event-related changes in the amplitude of the oscillatory neural response, data were collapsed across trials for left and right visual hemifield and the full-length artefacts-free epochs were used. We employed a complex Morlet wavelet analysis. We used 3 cycles at the lowest frequency and 16 cycles at the highest frequency. These parameters provide estimates of the event-related changes in the oscillatory power in 100 log-spaced frequencies from 3 Hz up to 80 Hz in a 200 time points ranging from −944 to 940 ms relative to the target onset. Baseline included all the time points before the target onset.

We aimed to test power changes especially in the beta (15-30 Hz) band, given the emerging role of beta oscillations in visual crowding (Ronconi *et al*., 2016*a*; Ronconi and Marotti, 2017; Battaglini *et al*., 2019). To assess the specificity of beta oscillation, we also tested alpha band (8-12 Hz) power changes, given its well-recognized role in visual processing and visual attention (for reviews see: Jensen *et al*., 2012, 2015; Klimesch, 2012). To detect reliable differences in these frequency bands, we applied N=10000 permutation tests with cluster-based correction for multiple comparisons performed in all scalp sensors (Maris and Oostenveld, 2007; Groppe *et al*., 2011) in the two frequency bands of interest, focusing on a time window after the onset of the target letter (0-600 ms). These tests were performed after collapsing data for the two crowding conditions (i.e. *strong* and *mid*), and after subtracting the activation in the *baseline* condition. This way we could estimate event-related oscillatory power changes which were specific to conditions requiring local visual processing in a crowding regime, independently from the activity related to the simple orientation discrimination of the target in isolation.

### Data analysis – Cortical source reconstruction

According to the most recent guidelines (Seeck *et al*., 2017; He *et al*., 2018), a reliable estimation of neural electrical activity at the cortical source level requires a scalp coverage with at least 64 EEG electrodes.

Since multiple brain regions are expected to be activated simultaneously in the time window of interests corresponding to significant ERPs variations, we opted for a distributed source imaging method, which is less user-dependent than fitting a restricted number of equivalent current dipoles. Forward modelling of neural electrical fields was computed through a Boundary Element Method (BEM) with OpenMEEG package (Gramfort *et al*., 2010), based on a realistic tessellation and conductivity profile of the head compartments: 1082 vertices for scalp, and 642 for outer and inner skull.

As a reasonable trade-off between complexity and completeness of the model, we chose: I) to decimate the default anatomical template into a grid of 15002 vertices, corresponding to a spacing of approximatively 5 mm between adjacent source locations on the cortical surface; II) to constrain position and orientation of the individual cortical dipoles orthogonally relative to the cortex, in order to obtain one electrical dipole at each of the 15002 vertices within the head volume.

The data segment between −500 and 0 ms relative to stimulus onset was used to estimate the noise covariance matrix. The estimation of distributed source amplitudes (in pA.m) was computed through a linear combination of EEG sensor time-series with a minimum-norm inverse kernel (MNE). To reduce the bias of the MNE toward superficial currents, a further depth-weighting step was introduced to adjust the current density estimates. A z-score normalization was applied to each cortical source trace with respect to the baseline period (−200, 0 ms): this standardization replaces the raw source amplitude value with a dimensionless statistic that is suitable for hypothesis testing. Such normalization reduces the influence of inter-individual fluctuations in neural current intensity that is due to anatomical or physiological differences of no (primary) relevance (Tadel *et al*., 2019).

Under the premise that the actual directionality of dipolar currents is of no interest for the present study, the absolute values were used to compute the contrast measure between two conditions (|A| - |B|) regardless to the currents polarity. Subject-level averages were obtained by weighting each condition-specific mean by the number of trials in the same condition.

After obtaining the individual (cortical) maps of source activity, a further surface smoothing was applied using a circularly symmetric gaussian kernel with a full width half maximum (FWHM) size of 5 mm. Such further step improves the possibility to detect differential activity in a specific cortical region at the group level by reducing noise and inter-individual variability.

At second-level analysis, the output of intra-subject means was averaged to produce the grand average of source data for each group (TD and ASD) and condition (*baseline*, *mid* and *strong* crowding levels) separately.

To better highlight the cortical activations generated by the increasing crowding task, activation in the *baseline* crowding condition was subtracted from the *mid* and *strong* crowding conditions (for a detailed discussion of the subadditive logic here applied at sensor and source space, see Stevenson *et al*., 2014).

### Data analysis - Functional connectivity

Connectivity analysis was performed in the source space rather than at sensor level to avoid the inaccurate assumption that specific sensor locations correspond across individuals, despite variable head shapes and orientations/positioning (Peelle *et al*., 2013). Furthermore, the extraction of cortical time series reduces the effect of electromagnetic field spread, and prevent spurious (non-independent) source-leakage effects, such as linear mixing or cross-talk between time series (Schoffelen and Gross, 2009).

To estimate the coupling between pairs of sources, we applied the phase-locking value (PLV), also called Mean Phase Coherence (MPC: Mormann *et al*., 2000) or inter-site phase clustering (ISPC: Cohen, 2015), which is a widely used class of measure for phase-dependent interactions (Schoffelen and Gross, 2009).

The PLV is a metric for quantifying absolute value of the mean phase synchronization between two narrow-band signals (Lachaux *et al*., 1999) under the hypothesis that connected (“locked”) areas generate signals whose instantaneous phase evolve together. Since the stimulus event resets the phase of the neural oscillators, two brain signals interacting at a given time should have a constant phase difference over repeated trials (Bruña *et al*., 2018). This reason makes the PLV a method particularly tailored to study evoked activity.

Differently from coherence measures, the PLV assumes that neural signals may synchronize their phases, without the necessity of simultaneous, increased power modulations: this make PLV robust to fluctuations in amplitude and less affected by the spurious influence of the (common) reference electrode (Nunez *et al*., 1997; Boersma *et al*., 2013).

Since two brain regions interact functionally by exchanging information in a specific narrowband, the first step in computing PLV is filtering the data in the frequency range of interest. For each trial, the wavelet transform was extracted on a time-window of 600 ms post-stimulus and then provided as input to the PLV computation. The measure of trial to trial variability for the (relative) phase of each seeds pair is expressed as a numerical value between 0 and 1: zero if there is no synchrony among the phases of the two signals and they are weakly connected, otherwise if the phases are strongly coupled in all trials the PLV will approach unity (Aydore *et al*., 2013).

A data-driven procedure was conducted to select from the entire brain the clusters of adjacent vertices that discriminate between the two groups of subjects in a specific level of crowding. The seed-to-seed connectivity matrix thus defined entered a group-level statistical test (independent-samples two-tailed *t* test) across the ASD and TD groups. A severe correction for multiple comparisons would determine the lack of statistical power for detecting any effect. For this reason we elaborated a non-parametric “regional”-based permutation method to assess the significance of our inter-group connectivity results, which is similar to the method proposed by Mamashli (Mamashli *et al*., 2019) for testing the difference between experimental conditions.

The procedural steps can be summarized as follows:

i. include *N* set of vertices (sub-regions of interest or sub-ROIs) in a functional coherent region (ROIs) and an approximately equal extent of *M* sub-ROIs in a different cortical area. See Table 2 for the list of all ROIs and the relative sub-ROIs included in the connectivity analysis;
ii. estimate for all the subjects and conditions the connectivity values between each pair of sub-ROIs, generating a *N x M* matrix of connectivity scores;
iii. compute an appropriate cluster statistic: in this case a two samples *t* test under the null hypothesis that the *N_(i)_M_(j)_* connectivity score (for each combination of sub-ROIs from the *N* and *M* clusters) have the same value in both TD and ASD groups;
iv. determine the cluster mass by summing the number of *N_(i)_M_(j)_* sub-ROIs pairs with a *t* value exceeding an a priori defined statistical threshold (|*t|* > 1.96 or *p* < 0.05). This number can be considered the observed regional connectivity value at the ROIs level;
v. permute *n* times (*n*=1000) the *N_(i)_M_(j)_* connectivity score at single subject level by randomly assigning (shuffling) each subject value to one class (TD) or the other (ASD);
vi. for each iteration, repeat step (iv) and (v) in order to derive the values of the (surrogate) null distribution with the number of *N_(i)_M_(j)_* pairs statistically significant by chance;
vii. if the observed regional connectivity value, i.e. the number of *N_(i)_M_(j)_* significative pairs, falls in the 95^th^ percentile (right tail) of the surrogate null distribution, then the connectivity between the two cluster regions composed by *N* and *M* subset of vertices is above the critical cut-off.

**Table 2.** Anatomical coordinates of the seeds included in the connectivity analysis are derived from the Desikan-Killiany atlas (as adopted in the Brainstorm software). For a more accurate reconstruction of the activations in the occipital lobe, areas and labels are extracted from the Destrieux parcellation in gyri and sulci.

| Index | MNI coord. | Anatomical Loc* | Function (Neurosynth) |
| --- | --- | --- | --- |
| 1 | (-1.1, 35.0 -12.4) | L cingulate anterior | decision making, choice, reward, social interactions |
| 2 | (-11.7, 44.0, -22.3) | L orbitofrontal lateral | regulation, discriminative |
| 3 | (-2.0, 32.8, -25.6) | L orbitofrontal medial | disgust, simulation, ASD, anger, valence |
| 4 | (-1.9, 64.6, -14.1) | L frontal pole | beliefs, emotional |
| 5 | (-47.6, 18.8, 35.8) | L frontal middle rostral | memory retrieval, recognition task, control process |
| 6 | (-46.3, 16.9, 33.1) | L frontal middle caudal | language, phonological, word, reading, retrieval, verbal, judgment, lexical |
| 7 | (-58.6, 29.3, 6.0) | L frontal inferior (pars triangularis) | sentence, comprehension, language network, word, reading, syntactic |
| 8 | (-52.4, 8.3, 6.1) | L frontal interior (pars opercularis) | motor, speech production, language network, imitation, articulatory |
| 9 | (-53.0, 7.5, 7.1) | L precentral | motor, language, naming, speech production, imitation, pseudowords |
| 10 | (-39.8, 19.9, -0.6) | L insula | ASD, pain, empathy, nociceptive |
| 11 | (-29.5, -6.6, -27.5) | L parahippocampal | faces, emotional, arousal, valence, ASD |
| 12 | (-34.5, 4.8, -50.6) | L fusiform | episodic, retrieved |
| 13 | (-30.6, 10.8, -45.3) | L temporal pole | recollection, morphology, mental state |
| 14 | (-45.5, 0.2, -39.2) | L temporal middle | social, motor network, mentalizing |
| 15 | (-54.9, -4.2, -37.6) | L temporal inferior | theory mind, social, memory, semantic, retrieval, nouns, reading |
| 16 | (-56.7, 14.8, -12.7) | L temporal superior | language comprehension, speech, words, sentence |
| 17 | (6.9, 33.7, -9.5) | R cingulate anterior | valence, emotional, reward |
| 18 | (2.2, 40.7, -20.3) | R orbitofrontal medial | theory mind, social interaction, mental state, ASD |
| 19 | (0.4, 63.6, -16.6) | R frontal pole | reward, impulsive, emotional, recognize |
| 20 | (18.6, 59.8, -20.9) | R frontal middle | heart rate, social, regulation |
| 21 | (19.6, 51.5, -20.4) | R orbitofrontal lateral | - |
| 22 | (16.2, -69.8, 26.0) | R cuneus (gyrus) | item, recollection |
| 23 | (22.2, -86.5, 18.7) | R occipital superior (gyrus) | vision, eye field, spatial |
| 24 | (21.3, -85.8, 20.4) | R occipital superior (sulcus) | visual, learn, eye field, choice |

A final important note should be stressed on how to interpret the results of such inferential process (Sassenhagen and Draschkow, 2019). The assumption behind the null distribution obtained with this procedure is that the cluster structure of the data is exchangeable between groups: no matter if the connectivity values were drawn from group 1 (TD) or group 2 (ASD), they have the same probability distribution.

Under these premises, connectivity results derived from the cluster-based permutation tests cannot be assigned to the specific elements (sub-ROIs) included in the cluster, but they should be confined at the cluster level. In other terms, this procedure allows to make inferences about differences in connectivity between ASD and TD only at the level of the whole functional ROIs but not at the finer grained scale of sub-ROIs (see Table). Such methodological approach, on the one hand, increases the statistical power by attenuating the multiple comparison issues that the traditional correction procedures (e.g. Bonferroni, FDR) face at the cost of an inflated type II error. On the other hand, the statistical inference derived from a set of widely spaced ROIs takes into account the limited spatial resolution of EEG source reconstruction thus avoiding incautious connectivity analysis performed on an overdetailed cortical parcellation.

## RESULTS

### Behavioral performance: comparisons between groups and intra-group correlations

In all experimental conditions both groups performed above chance (chance level=0.25%; mean accuracy rate ± SEM in the *strong* crowding ASD: 0.47 ± 0.08; TD: 0.51 ± 0.12; in the *mid* crowding ASD: 0.61 ± 0.14; TD: 0.64 ± 0.15; in *baseline* crowding ASD: 0.79 ± 0.12; TD 0.82 ± 0.11, Figure 1D). The ANOVA showed a significant main effect of the level of crowding (F_(2,72)_=288.37, p<.001), but no significant main effect of group (F_(2,72)_=0.028, p=.382) or interaction (F_(1,36)_=0.784, p=.972) emerged. We checked for potential difference in accuracy between stimuli presented in the left vs. right hemifield, but no significant effects emerged (*all ps>.557*).

We tested also if, despite the absence of group differences, there was a correlation at the individual level between accuracy in the crowding task and ASD symptomatology as measured by the SCQ score. We performed partial correlations where the effects of age and accuracy in the *baseline* condition were accounted for. Results showed that ASD symptomatology as measured by the SCQ questionnaire (current version) was positively correlated with accuracy in the crowding task for the *strong* crowding condition (r_(15)_= 0.462, *p*=.036; Figure 6A) and the *mid* crowding condition (r_(15)_= 0.529, *p*=.017).

### ERPs results

A repeated measure ANOVA carried out on P1 mean amplitude did not show a significant interaction between levels of crowding and groups (F_(2,72)_=2.868, p=.063). No main effect of crowding was found (F_(1,72)_=1.564; p=.216), nor a main effect of group (F_(1,36)_=0.500, p=.484). These results were confirmed when the same analysis was performed after subtracting the *baseline* from the two crowded conditions (i.e., *strong-baseline* and *mid-baseline*): the interaction between group and level of crowding was not significant (F_(1,36)_=2.956, p=.094), and there were no main effects of crowding level (F_(1,36)_=2.009;p=.165) and group (F_(1,36)_=2.765;p=.105).

The ANOVA on the N1 mean amplitude showed a significant effect of the level of crowding (F_(2,72)_=4.271, *p*=.018) but no significant main effect of group (F_(1,36)_=0.067, *p*=.797). Importantly, a significant interaction between level of crowding and group emerged (F_(2,72)_=3.554, *p*=.034; Figure 2C). Planned comparisons revealed that in the ASD group no significant differences between levels of crowding emerged (*all ps*>0.109). On the contrary, planned comparisons in the TD group showed that the N1 mean amplitude in the *strong* crowding condition was significantly different as compared to both the *mid* (t_(19)_=4.489, *p*<0.001) and the *baseline* crowding condition (t_(19)_=2.542, *p*=.02); no difference was found between *mid* crowding and *baseline* conditions (t_(19)_=-0.731, *p*=.474) (Figure 4a). These results suggested that only for TD children the N1 mean amplitude was significantly influenced by the degree of visual crowding. When contrasting the two groups with independent-sample t-tests, no difference between group emerged in any level of crowding (*all ps*>.379).

Crucially, we investigated the evoked neural activity specifically attributable to the effect of crowding (removing the activity evoked purely by the processing of an isolated stimulus) with an ANOVA performed after subtracting the activity elicited in the control condition (i.e., *strong-baseline* and *mid-baseline*). This analysis confirmed a significant main effect of crowding manipulation (F_(1,36)_=11.145, p=.002) and a significant interaction effect between level of crowding and group (F_(1,36)_=4.922, p=.033). The main effect of group was again not significant (F_(1,36)_=2.759, p=.105). Planned comparisons showed that there was a significant difference in the *strong-baseline* condition between the two groups (mean amplitude ± SEM in ASD group= −0.21±0.68µV, and in TD group = 0.45±0.78 µV; t_(36)_=-2.743, *p*= .009), whereas no significant difference was found in the *mid-baseline* condition (t_(36)_= −0.566, *p*= .575). When performing within-groups planned comparisons, it is worth noticing that while in the TD group a significant difference emerged between *strong-baseline* vs. *mid-baseline* conditions (t_(19)_= 4.49, *p* < .001), the same comparison did not result significant in the ASD group (t_(19)_= 0.71, *p=* .493). These latter results further suggest that although participants with ASD showed a different neural activity relative to their TD peers in the experimental conditions requiring the strongest local visual processing (i.e. *strong* crowding), they did not differentiate their neural activity between conditions with different degree of local visual processing (*strong* and *mid* crowding, after accounting for the *baseline* evoked response).

### Time-frequency results

In the alpha band (8-12 Hz) there was no significant difference between ASD and TD group in terms of power variations in the 0-600 ms post-stimulus time window (Figure 3C). On the contrary, we found significant differences in the beta band (15-30 Hz) in two separate clusters, one comprising occipital central channels and one comprising parietal and temporal left channels (cluster-corrected *p*=.041; Figure 3A, 3B). There was a tendency of significance also in a cluster of frontal left channels (cluster-corrected *p*=.072). Specifically, there was in all cases a beta power reduction after the stimulus onset for the TD group, while the ASD group showed an opposite trend with an increased event-related beta power. It is worth noticing that this analysis was performed after removing the activity of the *baseline* (uncrowded) condition, in which the target was presented in isolation. Thus, the difference in beta power found here was independent from the neural activity evoked during basic visual processing of the target stimuli and was instead specific for the local visual processing required to segregate the target letter from the flankers. The time-frequency plot in Figure 3B represents the activity of the significant channels, from which _i_t is evident that in the TD group there was a sustained beta power reduction after the stimulus onset. This desynchronization was not evident in the ASD group, which actually showed an opposite trend.

**Figure 3.**
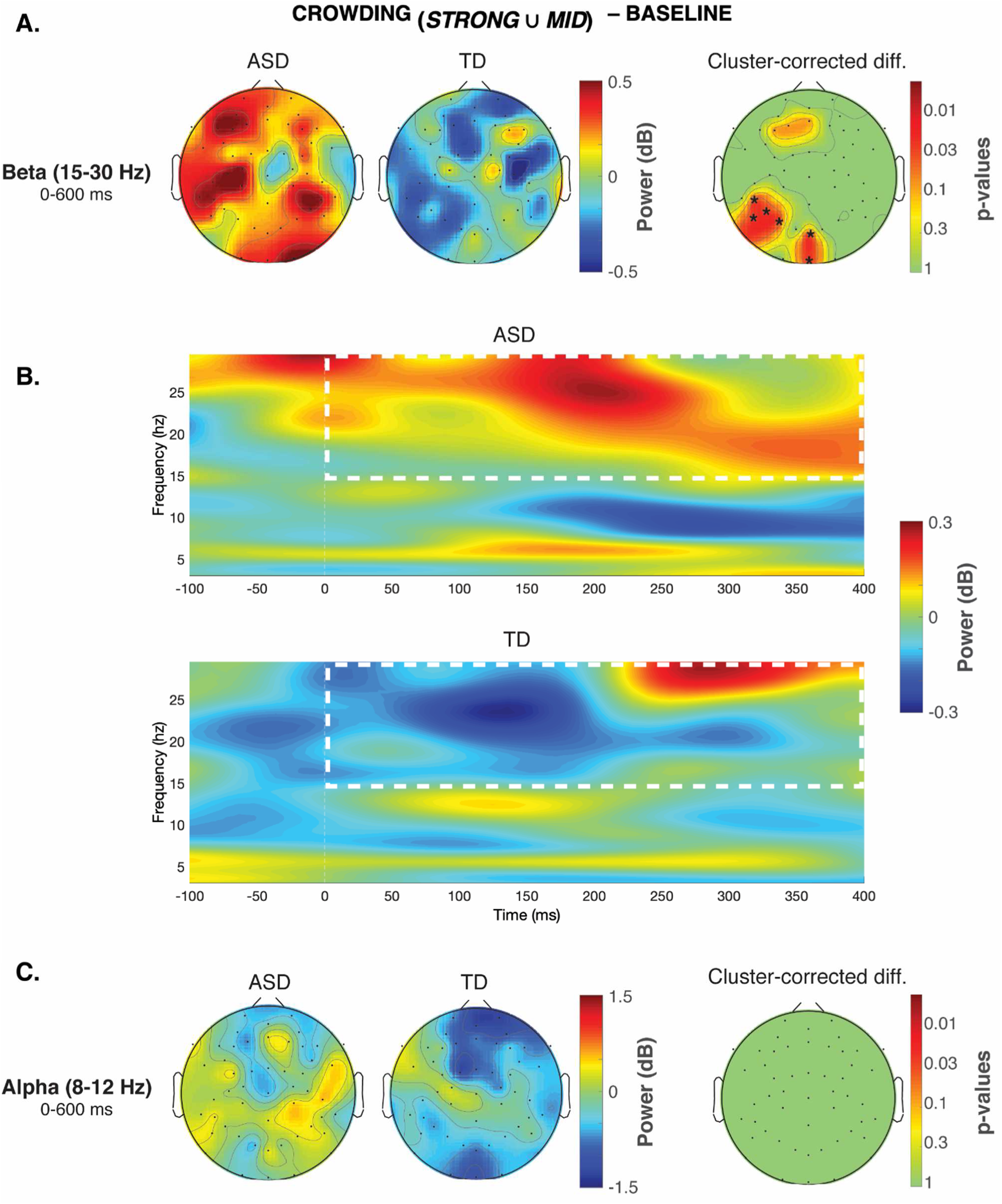
Event-related oscillatory activity reflecting detailed-oriented perception during visual crowding in ASD. (A) Scalp maps displaying the oscillatory amplitude (i.e. power) in the beta band (15-30 Hz) separately in the ASD and TD groups, together with the p-values map on the right side showing the significant cluster-corrected differences between groups (*=cluster-corrected p<.05). This analysis was performed after subtracting the oscillatory power in the *baseline* task condition (no flankers displayed), in order to highlight effects that emerged specifically within a crowding regime (*strong* and *mid*). (B) Time-frequency plot of the event-related oscillatory power averaged across all significant electrodes, showing a sustained decrement of beta power in the TD group after the target onset, which was not evident in the ASD group (that actually showed the opposite trend). (C) A similar analysis performed in the alpha band (8-12 Hz) did not reveal any significant differences between groups.

### Group-level source maps

Source estimates of neural activation are shown in Figure 4 for both TD and ASD group and for the time-windows of interest corresponding to the N1 (250-350 ms) ERP. The measure reported represents, in units of standard deviations (z-scores), how much the activity of a specific vertex deviates from the baseline period (−200–0 ms). All the activations reported have a minimum size of 5 vertices and overcome the statistical threshold (*p* < .05, uncorrected).

**Figure 4.**
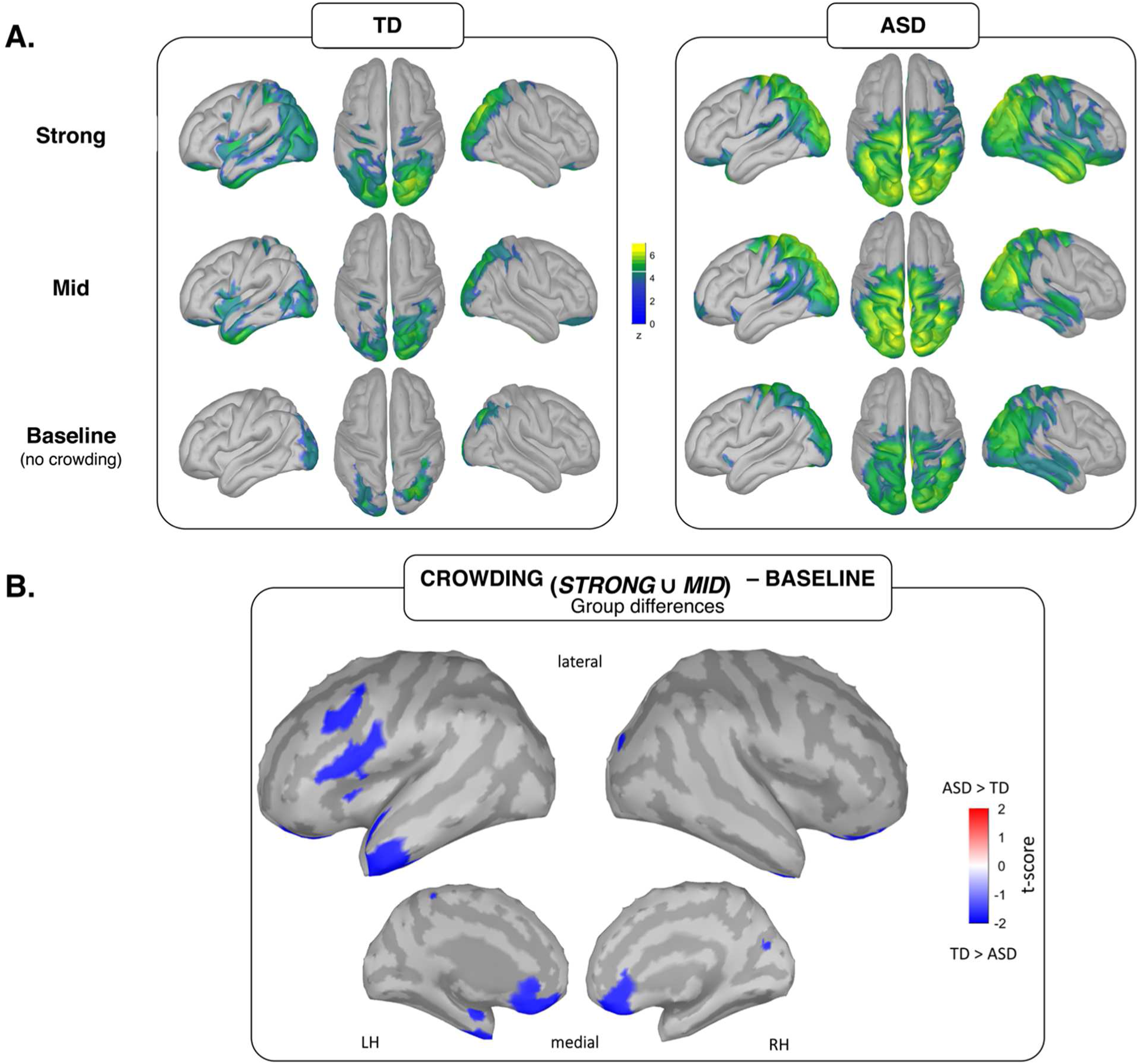
Cortical sources of visual crowding in TD and ASD groups. (A) Source estimates (z-scored values) of neural activity for each task condition (*strong* crowding, *mid* crowding and *baseline*) in the time-window corresponding to the N1 (250-350 ms) ERP. In the TD group, the pattern of activation recruits an increasing bilateral portion of the occipital, temporal and (superior) parietal cortex. Such modulation in the ASD group seems to be absent, especially when contrasting the mid and strong crowding levels. **(B)** Contrast maps between ASD and TD obtained after subtracting the neural response in the *baseline* condition. The cortical regions differently activated between the two groups spanned across the frontal and anterior-temporal portions of the left hemisphere, and the right occipital lobe (as listed in detail in Table 2: minimum size of 5 vertices, p<.05 uncorrected). In all these areas, the neural activation was significantly reduced in ASD as compared to the TD group.

In the TD group the activity was clearly modulated by the crowding levels, with a direct proportion between the target-to-flankers distance and the spread of activation: in the conditions where distractors were presented, the pattern of activation recruits an increasing bilateral portion of the occipital and (superior) parietal lobes.

In contrast, such modulation in the ASD sample was considerably reduced when comparing the control condition (*baseline*) with the two crowding conditions (*mid* and *strong*), and also when contrasting the *mid* and the *strong* condition. The enhanced responses obtained in the ASD group reveal that a comparable amount of neuronal resources in the occipital and temporo-parietal regions is allocated for processing the visual task independently from the crowding level. Overall, this pattern of activity mimics the absence of a significant N1 modulation in the ERPs.

As we did for the ERPs, to rule out the neural response to the target presented in isolation, we subtracted the activation pattern of the *baseline* from both the *mid* and *strong* crowding conditions, obtaining new maps of source activations that could be used to highlight the core neural areas showing differences in activations between the ASD and TD group that are specifically involved in segmenting the target from flankers. According to Gross et al. (Gross *et al*., 2013), t-tests at source level is only used to properly describe the cortical distribution of significant effects established at the sensor level: therefore, no correction for multiple comparison is required. The cortical regions that were differentially activated between the two groups as emerged from this analysis are reported in Figure 4B and listed in more details in Table 2, with MNI coordinates for each region, the matching structural brain locations as derived from the Desikan-Killiany atlas (Desikan *et al*., 2006) and keywords indicative of the regional functions retrieved from the *neurosynth* platform (Yarkoni *et al*., 2011). Specifically, we found different activity between groups in right occipital regions, left anterior temporal regions (spanning across superior, middle and inferior temporal gyri), left middle and inferior frontal regions, left anterior insula and bilateral orbitofrontal cortex. In all these regions activity was significantly reduced in the ASD group as compared to the TD group.

These cortical areas constituted the regions of interest for the connectivity analysis.

### Functional connectivity results

Based on the time-frequency results, we confined the investigation of the oscillatory components to the beta frequency range (15-30 Hz) in the cortical areas defined above (only areas exceeding the minimum threshold of 5 vertices were included). Doing so, the within-band coupling was computed on a total of 24 sub-ROIs.

Figure 5 displays the difference of connectivity in ASD against TD groups for the beta band conducted in crowding trials (*strong* and *mid*) once the *baseline* condition was subtracted. Each element in the PLV adjacency matrix represents an interaction between a pair of sub-ROIs and the colormap reports the value and the direction of the t-tests under the null hypothesis that the two seeds have the same PLV in both groups.

**Figure 5.**
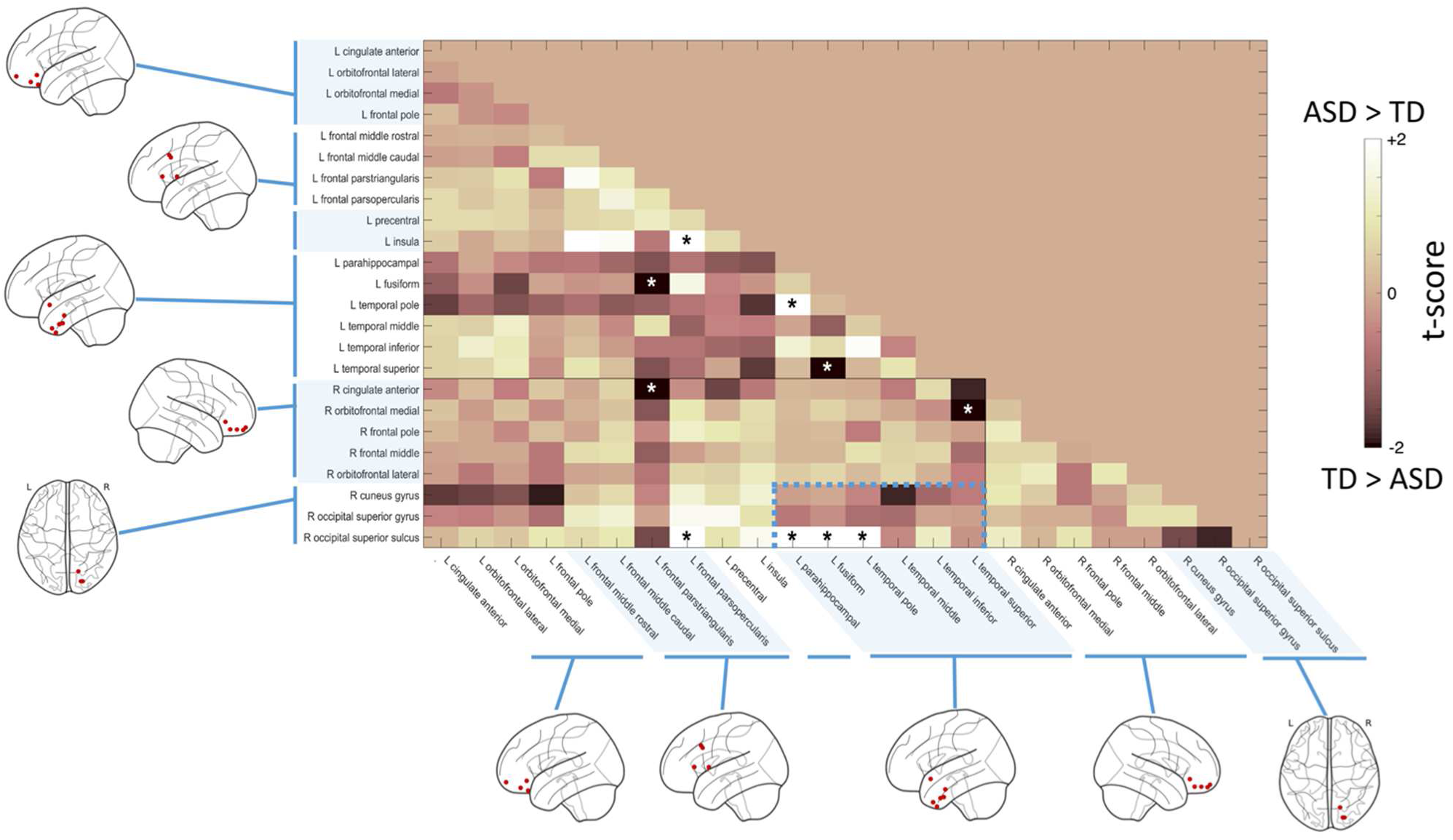
Functional connectivity in ASD vs. TD underlying detail-oriented perception during visual crowding. Phase locking values (PLV) were estimated for the 24 sub-ROIs identified/selected through the source reconstruction (Figure 4 and Table 2). Based on the time-frequency results, the connectivity analysis was conducted in the beta range (15-30 Hz). The connectivity matrix reports for each pair of seeds the value and direction of the statistical difference in connectivity between ASD and TD groups under a crowding regime. As assessed with “regional” cluster-corrected permutation tests, subjects with ASD showed an increased beta connectivity between the right occipital cortex and the left anterior-temporal lobe (p=.004; dashed line). The boundaries between hemispheres and anatomical regions (from left to right: orbitofrontal, prefrontal, temporal and occipital) are denoted by a solid line and different colour shades.

The statistical results assessed with the “regional”-based permutation procedure allows to make reliable inference at broad cluster level (ROI), and not for a single element of the cluster (sub-ROI). The pair of sub-ROIs showing significant cluster-corrected differences (*p*=.004) in connectivity between ASD and TD were located in the right superior occipital cortex (with activation spanning across the gyrus and sulcus, and some contribution of the cuneus), and the anterior portion of the left temporal lobe, in particular the fusiform gyrus.

The only connectivity pattern at regional level which survived the permutation analysis presented a uniform directionality of the effect: subjects with ASD showed a marked increase in the coupling between right occipital and anterior temporal ROIs as compared to the TD group, suggesting an abnormal reorganization of beta activity in a crowding regime.

No other significant differences in connectivity within or between hemispheres for beta frequency were found among groups.

Finally, although differences in oscillatory activity at the sensors level emerged only in the beta band, but not alpha band, we further checked whether these connectivity results were specific for the beta band by running the same analyses on PLV calculated in the alpha frequency range. We observed that no significant cluster-corrected differences in the alpha band emerged (*all ps*> .15).

### Correlations between beta-band connectivity, autistic symptomatology and detailed-oriented perception

We tested the possible relationship between the individual occipito-temporal functional connectivity index as measured with the PLV (by averaging the phase-locking values of the three pairs of seeds which survive the cluster-based permutation tests) and the ASD symptoms severity score as measured by the Social Communication Questionnaire (SCQ). Specifically, we performed partial correlations to account for individual differences in age. We found that in the ASD group there was a significant negative correlation between the individual average PLV index and the SCQ current version score (r_(15)_= −0.425, *p*=.045; Figure 6B). This correlation shows that individuals with the more severe ASD symptomatology where the ones presenting the more reduced beta-band connectivity between occipital and inferotemporal regions. No significant correlations between functional connectivity and SCQ scores emerged in the TD group (*all ps*> .176).

As a last analysis, we were interested in testing whether individual occipito-temporal functional connectivity was also related to behavioural measures of detail-oriented perception obtained during the visual crowding task. Again, a partial correlation analysis was performed to control for the effect of age and performance (accuracy rate) exhibited in the baseline condition. We observed a significant negative correlation between the individual functional connectivity scores (average PLV) and accuracy in the *strong* crowding condition (r_(15)_= −0.506, *p*=.045; Figure 6C); this result suggests that participants with ASD showing a diminished detail-oriented perception were the same exhibiting an increased beta-band connectivity.

**Figure 6.**
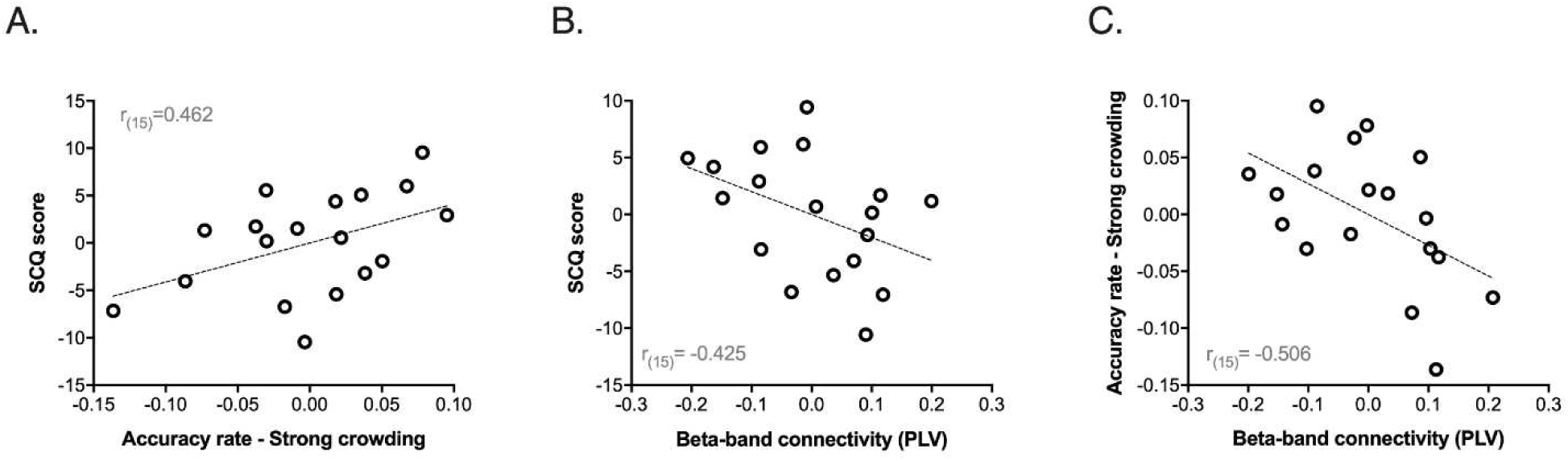
Correlations between ASD symptomatology, beta-band connectivity and behavioural performance during visual crowding. (A) Scatter plot showing the relationship between individual accuracy rate in the *strong* crowding condition and ASD symptomatology as measured by the Social Communication Questionnaire (SCQ) score (Current version) (a significant correlation emerged also for the *mid* crowding condition; see Results). (B) Scatter plot showing the relationship between individual occipital-temporal beta-band functional connectivity indexes (average PLV) and ASD symptomatology as measured by SCQ score. (C) Scatter plot showing the relationship between individual occipital-temporal beta-band connectivity and behavioural performance (i.e. accuracy rate) in the *strong* crowding condition. For the correlation analysis in (B), the effect of chronological age has been controlled for, whereas for the correlation analyses in (A) and (C) both the effect of chronological age and performance in the *baseline* task condition have been controlled for.

## Discussion

In the present study we employed a visual crowding task as a probe to test the neurophysiological correlates of detail-oriented local visual processing in individuals with ASD as compared to TD peers. Both groups showed a significant influence of crowding on their orientation discrimination accuracy, which became lower in conditions of stronger crowding. The finding of a comparable behavioural performance between the two groups is not surprising given the mixed results reported so far in the literature (Baldassi *et al*., 2009; Constable, 2010; Keita *et al*., 2010; Freyberg *et al*., 2016), and especially considering that in the present EEG study we employed only two target-to-flanker distances (together with a control condition with no flankers). This simplified EEG design with a limited number target-to-flankers distances could not have detected subtle behavioural differences in the threshold or slope of the psychometric function (see for example Battaglini *et al*., 2019). However, in agreement with previous studies reporting superior performance of individuals with ASD during visual crowding tasks (Baldassi *et al*., 2009; Keita *et al*., 2010), we found a significant positive correlation between accuracy in conditions of strong crowding and ASD symptomatology; specifically, in the ASD group participants with higher symptoms severity were those showing a better performance in the target-to-flankers discrimination task.

The confirmation of a different visual processing in a crowding regime came from the analysis of the N1 ERP component in occipito-parietal sensors. The mean amplitude of the N1 was significantly modulated in TD children; specifically, it was suppressed (i.e. lower amplitude) in a strong crowding regime where the target-to-flankers spacing was reduced. Such N1 suppression is in line with previous ERPs findings in the neurotypical adult population (Chicherov *et al*., 2014, Ronconi *et al*., 2016*a*) and also with other studies on texture segmentation, which typically found a significant N1 suppression when the segmentation of a target stimulus from the background was more demanding (Bach and Meigen, 1992, 1997; Caputo and Casco, 1999; Fahle *et al*., 2003). Contrarily to the results observed in TD children, such N1 modulation was absent in children with ASD that did not show any significant differences in this early ERP component as a function of visual crowding. When we accounted for the baseline neural activity evoked by the presentation of the target without flankers, we were able to highlight a significant difference in the ERP response between children with ASD and TD peers in conditions of strong visual crowding, confirming that although behavioural performance did not differ between the two groups, ASD children extracted local visual information differently in a crowding regime, over and above the difference that could be explained by the evoked neural response to the target presented in isolation.

We further analysed the evoked neural response and the group differences at the cortical sources level. Within the TD group, the cortical activity was modulated by visual crowding: an increase in crowding was accompanied by the spread of activation in the occipital, superior parietal and infero-temporal regions. Contrarily, such modulation of the spatial extent of activity at the source level was considerably reduced in the ASD sample: i.e. individuals with ASD showed a similar activation both when the target was presented in isolation and when it was surrounded by flankers. These results suggest that a comparable amount of neuronal resources in the occipital, parietal and infero-temporal regions was allocated during local visual processing independently from the amount of crowding, in line with the absence of a significant N1 modulation in the ERPs.

In terms of differences in activation between groups, after accounting for the baseline neural activity, we found a reduced activity in the ASD group as compared to the TD group in occipital regions, left anterior temporal regions, left middle and inferior frontal regions. The differences that we found in high-order visual areas in the inferior temporal regions were clearly localized in the left hemisphere, in agreement with neuroimaging evidence showing a pronounced left lateralization of the ventral stream for processing for processing single letters, groups of letters and words (Dehaene *et al*., 2005; Vinckier *et al*., 2007).

As a further step in the characterization of the neurophysiological correlates of local processing in ASD, we tested event-related power changes in beta band oscillations at the sensor level. Invasive recordings in non-human primates and M/EEG studies in humans consistently showed that variation in oscillatory activity in alpha and beta bands reflects reentrant feedback loops and large-scale cortical interactions (Donner and Siegel, 2011; Bastos *et al*., 2015; Jensen *et al*., 2015; Zaretskaya and Bartels, 2015). As reviewed in detail in the Introduction, such reentrant feedback information carried by beta-band neural oscillations might be a relevant mechanism for resolving visual crowding. Accordingly, previous evidence from our group showed that beta power was selectively associated to better crowding resilience (Ronconi *et al*., 2016*a*; Ronconi and Marotti, 2017), and non-invasive electrical stimulation (i.e. tACS) in the beta band leads to a beneficial effect on crowded vision, supporting a causal link (Battaglini *et al*., 2019). In the present study, we further replicated this evidence in our control sample of TD children, by showing an event-related desynchronization (i.e. power reduction) in the beta band spanning both pre- and post-stimulus time windows, after controlling for the baseline neural activity evoked by the target presentation without flankers. This beta power reduction was not evident in the ASD group, which differed significantly in the amplitude of the event-related beta activity from their TD peers in central occipital and left temporo-parietal channels. We also tested potential differences in the oscillatory power within the alpha band. Although alpha has not been directly associated to visual crowding (Ronconi *et al*., 2016*b*), it is largely recognized to have an important role in visual perception (for reviews see Jensen *et al*., 2012; Klimesch, 2012; VanRullen, 2016). However, no differences between groups emerged in this band, confirming previous findings of a specific association between beta oscillations and visual crowding and, importantly, suggesting that the ASD group did not show a general alteration of oscillatory response in detail-oriented tasks, but a specific alteration of top-down reentrant feedback signals carried on selectively by beta oscillations.

To investigate more deeply the hypothesis that the alteration of beta oscillations revealed in the ASD group at the sensors level reflected an alteration of top-down reentrant feedback connectivity coming from higher-order visual areas down to early visual regions, we performed phase-based (i.e. PLV) functional connectivity analyses at the cortical sources level in the beta band. In particular, we used seeds obtained by contrasting neural activity between ASD and TD in response to visual crowding. Cluster-corrected group differences in inter-regional connectivity emerged between early occipital and anterior infero-temporal areas. These results on the one hand confirm the hypothesis that beta oscillations reflect large-scale cortical interactions and, importantly, brings new evidence about the precise neurophysiological nature of enhanced local visual processing in ASD. Specifically, the present functional connectivity result shows that ASD were characterized by a systematic increase in connectivity between occipital and infero-temporal regions as compared to their TD peers. When testing the association between this altered functional connectivity pattern and autistic symptomatology, we found a significant negative correlation: ASD individuals showing the higher beta-band connectivity were those with the less severe symptomatology. Moreover, correlational analyses revealed also a significant negative correlation between occipito-temporal functional connectivity in the beta band and behavioural measures of detailed-oriented perception within a crowding regime. It is important to underline that such results were obtained after controlling for behavioural performance in the *baseline* (uncrowded) condition. From this combined evidence, a candidate hypothesis is that EEG functional hyper-connectivity in the beta band could reflect a possible compensatory mechanism recruited in particularly demanding crowding regimes that allows children with ASD to perform a balanced analysis of the visual scene where local visual information is weighted in the more global analysis of the visual scene. In the context of the present study this would lead to a diminished detail-oriented perception and a decreased discrimination accuracy within the crowding regime, possible indexing a more pronounced susceptibility to the disturbing effect of flankers.

Our findings of hyper-connectivity find support in recent studies testing whole-brain connectivity in ASD (Keown *et al*., 2013; Rudie and Dapretto, 2013; Supekar *et al*., 2013). Keown et al. (2013) used resting-state fMRI (rs-fMRI) to compare maps between adolescents with and without ASD and reported an anterior-posterior gradient of local under-to over-connectivity in ASD. In particular, occipitotemporal regions showed diffuse overconnectivity in ASD. Supekar et al. (2013) used rs-fMRI and a systematic whole-brain connectivity approach to analyze intrinsic brain connectivity in younger children with ASD; their results, which were replicated in two additional independent cohorts, showed that connectivity was diffusely increased in ASD regardless of physical distance, i.e. both short- and long-range connections were stronger in ASD.

Overall, these previous functional neuroimaging results agree with our EEG evidence in suggesting that increased connectivity in occipito-temporal regions may be related to islets of superior functioning in sensory systems of individuals with ASD (Rudie and Dapretto, 2013).

Regarding M/EEG evidence of altered functional connectivity in ASD, studies have primarily focused on the high frequency gamma band, reporting mixed findings (for review see Vissers *et al*., 2012; O’Reilly *et al*., 2017). Recently, Seymour et al. (Seymour *et al*., 2019) found differences between ASD and TD groups in the feedback connectivity between visual areas V4 and V1 mediated by alpha oscillations, with reduced modulation during simple visual presentation. In another MEG study looking at functional connectivity in ASD, Buard et al. (Buard *et al*., 2013) reported higher beta and gamma band functional connectivity specifically in the left hemisphere in the ASD group relative to controls between occipital and temporal regions (i.e. superior temporal gyrus). It is interesting to note that evidence of increased M/EEG functional connectivity in the beta band has also been reported during processing of somatosensory stimuli (i.e. passive tactile stimulation) in a study by Khan et al. (Khan *et al*., 2018).

Our multi-level approach to local visual processing in ASD with a well-controlled paradigm raised a series of promising clinical insights. This study further confirms the need of decomposing complex constructs (e.g., sensory and perceptual processing) into more refined and defined building blocks (e.g., local visual processing) in order to account for the heterogeneity of ASD, especially when using sophisticated analyses on time-resolved M/EEG signal. Beyond the idea that ASD is just a constellation of behavioral symptoms, recent experimental and theoretical perspectives converge in stressing “inflexibility”, “rigidity”, “habit of sameness”, and “absence of modulation” as key (and core) features of ASD. This is reflected in clinical and computational perspectives underlying inflexibility of learning, thinking and acting (D’Cruz *et al*., 2016; Lawson *et al*., 2017; Casartelli *et al*., 2018). Intriguingly, this general framework nicely fits with the present findings: despite a comparable behavioural performance, TD and ASD groups differed in their event-related neural activity at the sensors and sources level. In TD participants, neurophysiological results mirrored behavioral results by showing a clear modulatory effect of the crowding levels. Such adaptive (flexible) neural processing of different visual scenes observed in the typical processing is almost absent in the ASD group: in contrast, their neural processing is rigid (inflexible) and prone to process all visual scenes in a similar way. Such dissociation between flexible (TD) and inflexible (ASD) balancing may represent a promising paradigm shift in the context of ASD research (Casartelli, 2019).

A critical challenge for studies exploring basic mechanisms as putative building blocks for more complex functions concerns the link with behavioral outcomes/symptomatology. Recently, Wang and colleagues (Wang *et al*., 2015) used a machine learning approach to analyse eye-tracking data of individuals with ASD looking at different common visual scenes (e.g. pictures of people on the beach or depicting soccer/football matches). By adapting a three-layered saliency model that took into account distinct attributes (pixel-level; object-level; semantic-level), the authors suggested that individuals with ASD *did not see* the “same” scene as TD individuals. This might in turn explain their different comprehension of the pictures and their different interpretation of the visual environment. Following this hypothesis, atypical behavioural responses that are often considered the core feature of ASD may “simply” be coherent with their specific interpretation of the visual scene. We do not intend to claim that anomalies in visual processing can explain all clinical manifestations of ASD, neither that visual processing anomalies have a sort of prominence as compared to other candidate markers of ASD. Nevertheless, state-of-the-art literature on local visual processing and our own study strongly suggest that heterogeneous clinical manifestations of ASD may be at least partially explained by anomalies in processing and encoding basic features of the visual scenes.

To conclude, visual processing in crowding regimes represents a well-established probe to explore detail-oriented visual processing in ASD. Starting from the widely recognized critical role of sensory and perceptual anomalies in ASD, we provide the first neurophysiological characterization of their detail-oriented visual processing style which has been largely investigated at the behavioural level because of its potential impact on higher-order functions connected to emotion recognition, speech processing and social interactions (Wang *et al*., 2015; Jarvinen-Pasley *et al*., 2008; Vlamings *et al*., 2010). Future challenges in the context of ASD research concern the possibility of benefiting from this body of evidence to address specific function-(e.g., visual processing) and component-(e.g., local visual processing) based rehabilitative approaches, in order to promote valuable cascade effects on more complex abilities. How the characterization of sensory and perceptual processing derailments from the typical developmental trajectory in ASD may be useful and feasible to this purpose is a critical issue for clinicians (Gliga et al., 2015). Clarifying the precise neurophysiological counterparts of these functions and their putative anomalies in ASD is a mandatory preliminary starting point.

## Acknowledgments

This work was supported by the “5per1000” funds for biomedical research (Scientific Institute IRCCS Medea; 2016, 2017 to LC), and by the Italian Ministry of Health (RC 02016-2018 to LC). The funders did not participate in the conception and development of this work.

The contribution of the clinical staff of the Scientific Institute IRCCS Medea as well as of children and their families are gratefully acknowledged.

